# Generative Reconstruction of Unobserved Cellular Dynamics using Single-Cell Transcriptomic Trajectories

**DOI:** 10.64898/2025.12.29.696948

**Authors:** Sumanta Ray, Sukhen Das Mandal, Snehalika Lall, Saumyadipta Pyne

## Abstract

The primary challenge in cellular dynamics is to capture the intermediate states between two distinct biological stages. This is because of the technical constraints in capturing transient states or the nature of single-cell sequencing protocols. To address this challenge, we introduce a new computational framework, GRAIL (**G**enerative **R**econstruction of **A**rtificial **I**ntermediate **L**ineages), which aims to bridge these missing transitions. GRAIL reconstructs biologically plausible intermediate cell states between two distinct developmental stages (A and B) through a Locality Sensitive Hashing (LSH) based cell selection strategy followed by a smooth interpolation in learned latent space. The interpolated latent representations are subsequently decoded into gene expression profiles using a trained generator from a Generative Adversarial Network (GAN). The framework consists of three components: (a) a pretrained autoencoder that learns latent representations of stage specific transcriptomes, (b) an LSH-guided interpolation procedure that identifies anchor cells and performed interpolation in latent space, and (c) a GAN generator that extrapolates realistic intermediate expression from the interpolated samples consistent with the underlying trajectory. We benchmarked GRAIL with different state-of-the-arts (SOTA) in diverse setup of simulated dataets as well as in real life scRNA-seq datasets. Generated samples from GRAIL preserve expected marker gene expression patterns and also improve downstream analyses, including differential expression and cell clustering. Our method addresses the critical gap in studying cellular transitions when experimental intermediate samples are unavailable.

## Introduction

Single-cell transcriptomic analyses have enabled researchers to reconstruct cellular trajectories and thus capture dynamic changes in gene expression during processes such as differentiation, regeneration, and disease progression. By ordering cells along a pseudotime axis, trajectory inference reveals the temporal evolution and branching of cell states, in contrast to static clustering that only identifies discrete cell types. This approach has proven transformative, offering unprecedented precision and depth in understanding complex biological systems. It allows identification of lineage-specific genes and pathways associated with particular trajectory branches, yielding insights into the molecular drivers of cell fate decisions and developmental progressions.

Computational methods for trajectory inference have increased rapidly in recent years. Classical pseudotime algorithms such as Monocle^1^ first introduced an unsupervised strategy of ordering single cells in pseudotime by learning an explicit principal graph that connects cells along a continuous developmental path. It laid the groundwork for delineating smooth gene expression changes accompanying cell state transitions. Other methods like Slingshot^2^ infer multiple branching lineages by fitting smooth curves through clusters of cells, enabling robust pseudotime estimation even in complex bifurcating trajectories. Graph-based approaches such as PAGA (partition-based graph abstraction)^3,4^ constructs an interpretable graph of cell clusters based on their transcriptomic connectivity, preserving the global topology of continuous transitions in the data manifold. These and related tools (e.g. diffusion pseudotime and graph kernels) have become popular for generating developmental trajectories from single-cell RNA-seq (scRNA-seq) data^5–8^.

In parallel, more advanced frameworks have leveraged additional information and deep generative modeling to improve trajectory reconstruction. For example, RNA velocity^9^ provides a temporal derivative for each cell by comparing spliced and unspliced mRNA, thereby inferring the future direction of cell state changes. scVelo^10^ extend this idea with a dynamical model that solves transcriptional kinetics by generalizing RNA velocity to transient cell states and recovering each cell’s position along an underlying differentiation timeline^11^. Other methods explicitly model cell state distributions over time. Waddington-OT (WOT)^12^ utilized optimal transport theory to infer temporal couplings between cell populations at different time points. It treats snapshots as probability distributions in gene expression space and computes the optimal flow of cells from an initial to a later state. TrajectoryNet^13^ builds an optimal transport which links continuous normalizing flows with dynamic OT to allow nonlinear, continuous-time mappings between distributions. This deep learning approach constrains a neural ordinary differential equation (neural ODE) with an OT-based loss, yielding smoother and more realistic trajectories than static OT interpolation alone. Deep generative architectures have also been explored for trajectory tasks. For instance, the Velocity Autoencoder (VeloAE)^14^ uses a tailored autoencoder to denoise high-dimensional expression data and learn a low-dimensional embedding of RNA velocities, enabling more robust detection of cell state transitions despite technical noise. Hybrid strategies (e.g. scPath^15^ along with its energy-based model scEnergy) attempt to map cells onto a putative developmental energy landscape, conceptually similar to entropy-based methods like SCENT^16^, to infer paths of differentiation down a potential gradient. Together, these methods represent a growing trend toward using deep learning and probabilistic modeling to capture complex, nonlinear trajectory structures that classical algorithms might miss.

Despite this progress, trajectory inference in single-cell data faces several key challenges: Experimental protocols often capture only endpoints or sparse time points, making it difficult to observe transient intermediate cell states. Additionally, datasets compiled from multiple sources can exhibit batch-specific biases, complicating trajectory reconstruction^17,18^. As a result, pseudotime ordering or cell transition estimates can be unstable or inaccurate. Effective denoising or dimensionality reduction is often required to discriminate true biological signal from technical noise^19,20^. Many biological processes involve branching trajectories (e.g. multipotent stem cells differentiating into several lineages) or other nonlinear progressions. Inferring these branching structures remains difficult. Moreover, many biological processes involve complex branching or nonlinear trajectories which are difficult to identify.

Existing advanced methods address some of these issues only partially. For example, Waddington-OT^12^ can model cell proliferation or death by allowing unbalanced transport (adding or removing “mass”)^21^, and it provides a smooth coupling between distributions rather than treating time points independently. TrajectoryNet^13^ similarly enforces smoothness and continuity by design, avoiding implausible jumps in state transitions^22,23^. However, these approaches rely on assumptions of gradual change and require either multiple intermediate time points or additional information (e.g. RNA velocity) to constrain the dynamics. Neither OT-based couplings nor continuous normalizing flows inherently generate new gene expression profiles, instead they interpolate existing ones or push points through a learned vector field. As a result, their intermediate states may lack realistic variation, especially in under-sampled regimes. In sparsely sampled or endpoint-only datasets (where only an initial and final state are profiled), such models struggle to recover complex trajectories, and they may default to overly smooth or linear transitions and miss rare branching events. There is a need for a method that can extrapolate plausible intermediate cell states with minimal assumptions, even when data are limited to extreme time points.

Here, we introduce GRAIL, a generative adversarial network-based framework for single-cell trajectory inference, to overcome these limitations. GRAIL combines latent space interpolation with adversarial data generation to model developmental progressions with improved realism and robustness. In our approach, a pretrained autoencoder^24^ is first used to embed high-dimensional scRNA-seq data into a latent space that captures the underlying biological structure. Within this latent space, GRAIL performs interpolation between source and target cell populations (e.g. an initial and terminal cell state), effectively constructing a continuous trajectory in a reduced dimension. Crucially, we leverage representative “anchor” cells from the starting state to guide this interpolation. The latent vectors of these anchors are perturbed and gradually moved toward the latent representation of the ending state, tracing a path through latent space. A pretrained GAN generator^25–28^ is then used to map these interpolated latent points to synthetic gene expression profiles, i.e. to generate realistic intermediate cell states along the trajectory. By incorporating stage-specific information in the GAN (for instance, conditioning on the identity of the starting vs. ending stage), GRAIL ensures that generated cells reflect the appropriate transcriptional progression. This design interpolating in a denoised latent manifold and using an adversarial network to output gene-level data that allows GRAIL to create transitional cell profiles that are both biologically plausible and realistic to the endpoints identity.

Innovations of GRAIL include its ability to bridge very distant cell states through a smooth latent interpolation, something that would be difficult in the original gene space due to noise and sparsity, and its use of a GAN to introduce variability and realism beyond simple averaging of transcriptomes. This results in markedly improved performance on challenging scRNA-seq trajectory inference tasks. In simulations where ground-truth intermediate states are known, GRAIL’s generated cells align much more closely to the true trajectories than those from previous methods, for example, yielding substantially lower Wasserstein distances between generated and real intermediate distributions (indicating better global gene-expression fidelity) and higher correlation with true pseudotime ordering.

GRAIL represents a new paradigm for single-cell trajectory inference, uniting deep generative modeling with pseudotemporal analysis to overcome the limitations of prior methods. It enables researchers to reconstruct realistic intermediate cell states and continuous gene expression trajectories from limited or static snapshots of data. This advancement opens the door to investigating dynamic biological processes (developmental lineage decisions, regenerative processes, tumor evolution, etc.) in contexts where traditional time-course sampling is impractical.

## Results and Discussions

### GRAIL Workflow

Figure 1 describes the workflow of the analysis pipeline that we have followed to get intermediate states within start state (stage A) and end state (stage B). GRAIL’s trajectory inference pipeline consists of four major components: (i) selecting representative anchor cells, (ii) encoding those cells into a latent space, (iii) interpolating latent representations, and (iv) generating high-dimensional gene expression data from the interpolated points. Figure 1 provides an overview of this workflow, with Figure 1, panel-A illustrating the anchor selection and encoding steps, panel-B showing the latent interpolation and final data generation steps. In the following sub-sections, we detail each component of the pipeline.

**Figure 1.**
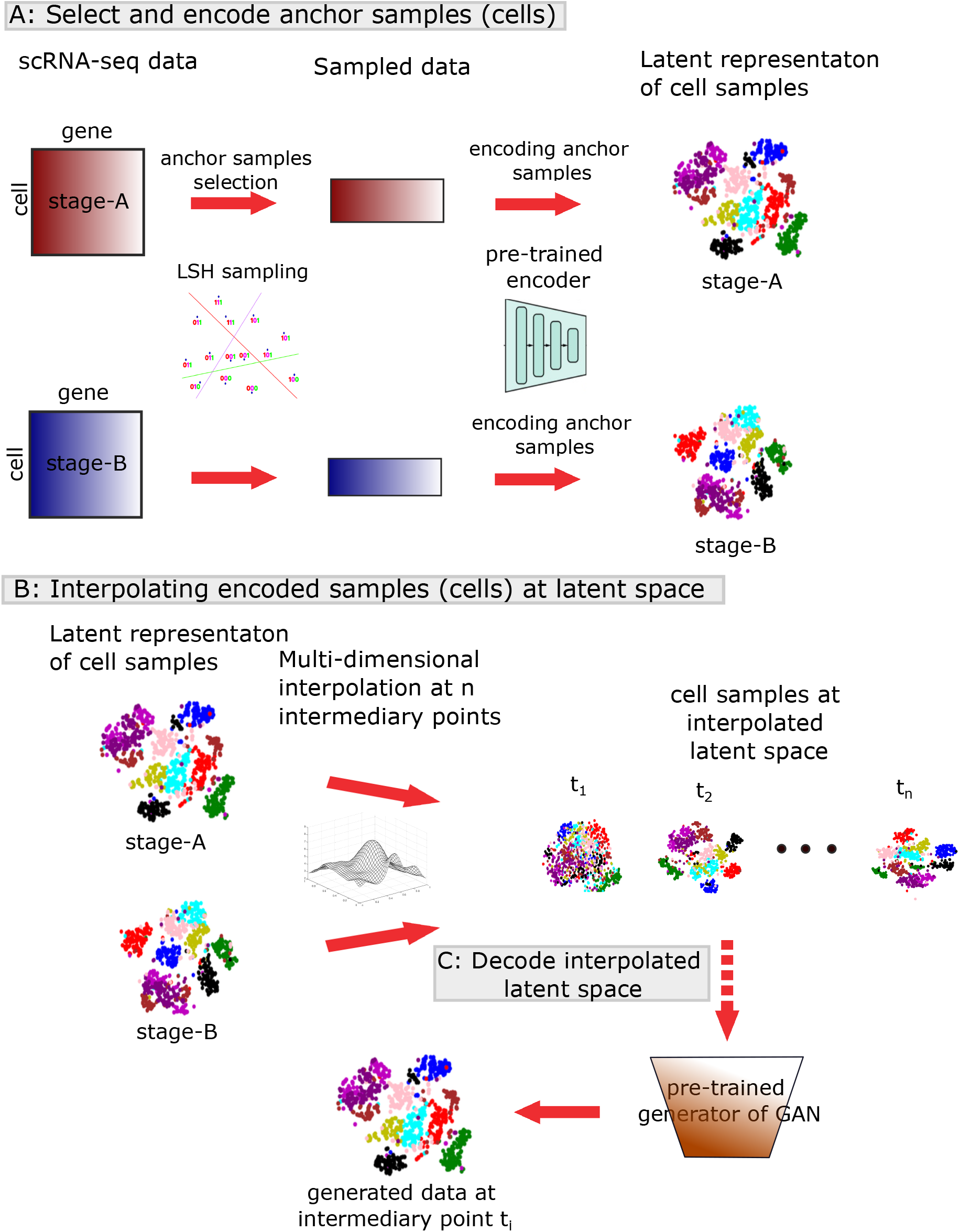
Overall workflow of the proposed model. Panel A illustrates the procedure for obtaining the latent representations of samples from each stage using Locality Sensitive Hashing (LSH) and an encoding process. Panel B depicts the generation of intermediary samples from the latent representations of each stage.

#### Anchor Selection

The first step in the GRAIL method is to identify a subset of representative cells—termed *anchorcells*—from both the starting population (stage A) and the ending population (stage B). We employ Locality Sensitive Hashing (LSH) to efficiently select these anchors (Figure 1, panel-A). LSH hashes cells with similar transcriptomic profiles into the same buckets with high probability, allowing us to sample diverse exemplars from each bucket. By drawing anchor cells from across different hash buckets (i.e., different regions of the gene expression space), we ensure maximal diversity and coverage of each stage’s expression landscape. This strategy preserves the salient features and the complex distribution of the original scRNA-seq data in a much smaller set of cells. The anchor cells from stage A and stage B serve as key reference points for downstream analysis: they capture the biological variation at the two endpoints and will guide the construction of the intermediate trajectory.

#### Latent Representation via Autoencoder

After selecting anchor cells, GRAIL embeds these high-dimensional gene expression profiles into a low-dimensional latent space using a deep autoencoder (Figure 1, panel-A). We train the autoencoder on the combined anchor datasets from stage A and stage B (see Figure 2, panel-A), so that the learned latent space encompasses the transcriptional variability of both endpoints. The autoencoder model has an encoder–decoder architecture: it compresses input gene expression vectors into a latent representation and then reconstructs the input from this code. Training is performed by minimizing the reconstruction error on the combined set of stage A and B anchor cells. This joint optimization ensures that the encoder captures features common to both states. Once training converges, we discard the decoder and retain only the encoder network for subsequent use.

**Figure 2.**
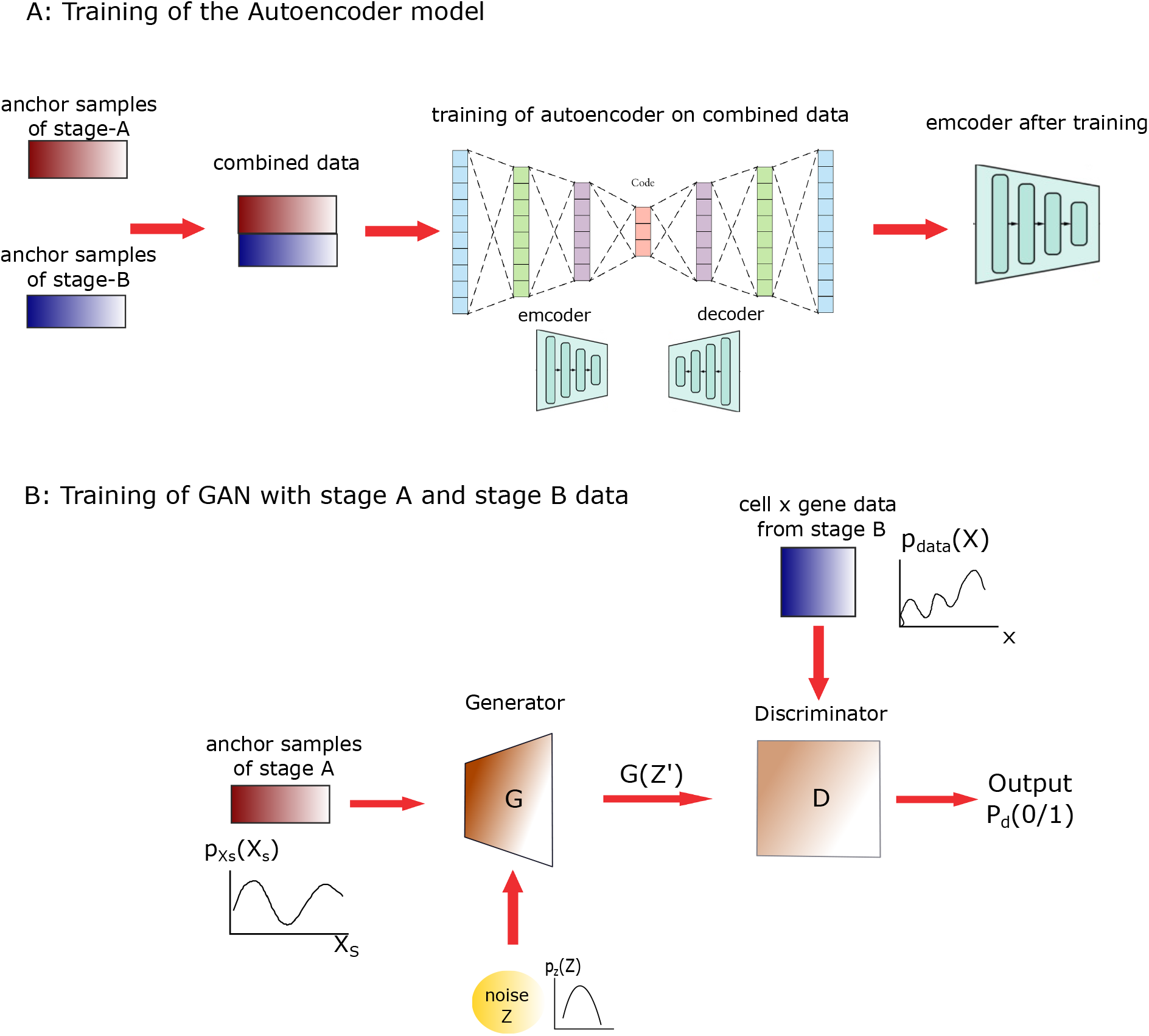
Training of the Autoencoder model and GAN. Panel A illustrates the procedure for training the autoencoder using the combined data from stage A and stage B. Panel B describes the training of the GAN model using anchor samples obtained from stage A and stage B

The trained encoder is then applied to all anchor cells to obtain their latent representations. As a result, each anchor cell (from both A and B) is represented by a vector in the shared latent space. Notably, this latent space provides a denoised, continuous manifold of cellular states that is well-suited for interpolation. Because it was learned from both stages jointly, the encoder’s latent space preserves the global structure of the developmental progression, making it possible to interpolate smoothly between stage A and stage B in subsequent steps.

#### Interpolation in the latent space

With the anchor cells encoded in latent space, the next step is to generate intermediate latent vectors that represent hypothetical cell states between the two original stages. We perform linear interpolation in the latent space between the representations of stage A and stage B (Figure 1, panel-B). In practice, for a given trajectory, we take latent vector of an anchor cell from stage A and a corresponding reference latent vector from stage B (for example, an anchor from stage B or the centroid of stage B’s latent distribution) and interpolate between them. A series of points is sampled along the line segment connecting the start and end latent vectors, producing a sequence of latent states at progressive intermediate stages *t*_1_, … *t*_*n*_ (where *n* is the desired number of steps between A and B).

Each interpolated latent vector thus represents a putative cell state at some pseudotime between the initial and terminal populations. Because the interpolation is done in a compressed, biologically informed latent space, these intermediate representations are expected to vary smoothly and plausibly, avoiding the noise and sparsity that would complicate interpolation in the original gene expression space. As illustrated in Figure 1, panel-B, this process produces a continuum of latent points bridging stage A to stage B. In the final step of the pipeline, these latent points will be mapped back to high-dimensional gene expression space using a generative model.

#### GAN-Based Generation

To reconstruct realistic gene expression profiles from the interpolated latent vectors, GRAIL leverages a Generative Adversarial Network (GAN). We design and train a GAN model to learn the mapping from latent space to the distribution of stage B gene expression, while incorporating information from stage A in the input. The GAN consists of a generator network *G* and a discriminator network *D*. During training (see Figure 2, panel-B), the generator takes as input a composite vector that includes an encoded stage A anchor cell along with a random noise component. In other words, instead of feeding only uniform noise as in a conventional GAN, we provide *G* with a vector 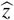 that combines features of a stage A cell (captured by its latent encoding) and stochastic noise. The generator 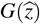 then produces a synthetic gene expression profile. The discriminator *D* is presented with two kinds of samples: (i) real cells from stage B, and (ii) generated outputs 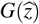 (where 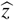 is sampled from the distribution of stage A anchors combined with noise).

So, the value function can be written as:

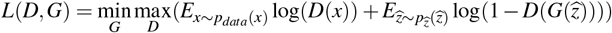

D and G form a two-player minimax game with value function L(G,D). We train D to maximize the probability of correctly validating the data of stage B and generated data. We simultaneously train G to minimize 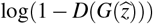, where 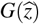 represents the generated data from the generator by taking the noise (pz) and the sample data of stage A 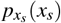 as input. As generator function takes sample data of initial stage A, so trained generator stores the information of transition from stage A to final stage B.

After this GAN training phase, we retain the generator as a powerful decoder to translate latent vectors into realistic gene expression profiles. We then use the trained generator *G* to map each interpolated latent point (from the latent interpolation step) into the high-dimensional gene expression space (Figure 1 panel-B). Because the inputs to *G* incorporate stage A-derived features and lie along a path toward stage B in latent space, the outputs are synthetic cells that represent progressively intermediate states between stage A and stage B. As the interpolation progresses from the starting anchor’s latent vector toward the ending latent vector, the generator produces a sequence of cells that traverses the transcriptional trajectory from the initial to the terminal stage. This combination of latent-space interpolation and adversarial generation ensures that the resulting intermediate cell profiles are both smoothly varying (reflecting a continuous transition) and biologically plausible (resembling real cells at or between the endpoint stages). Together, these steps constitute the GRAIL workflow for reconstructing developmental trajectories from endpoint-only scRNA-seq data.

### Performance evaluation of GRAIL on diverse simulated setup

#### Generation of simulated data

To rigorously evaluate the effectiveness and generalizability of our proposed framework, we generated a diverse set of simulated scRNA-seq datasets using the Splatter simulation toolkit. Splatter provides a flexible platform for simulating realistic single-cell RNA-seq data under various biological and technical conditions, allowing us to systematically test our method across a broad spectrum of scenarios.

Table 1 summarizes the key characteristics of the seven synthetic datasets (P1–P7) used in our evaluation. Each dataset models a pair of cellular stages (Stage A and Stage B) with controlled properties such as population balance, differential gene expression (DE) levels, dropout rates, and trajectory topologies. These configurations enable comprehensive benchmarking of our method’s ability to interpolate intermediate states under both ideal and challenging conditions. Dataset P1 simulates a balanced linear transition with moderate dropout and 40% of genes exhibiting differential expression across three distinct cell types. P2 introduces severe population imbalance (100 cells in stage A vs. 800 in stage B), mimicking real-world scenarios such as rare-cell detection and enrichment-based sampling biases. P3 models subtle gene expression changes (low DE proportion of 10%) to assess our method’s sensitivity in detecting gradual and nuanced cellular transitions. P4, by contrast, simulates strong expression changes with high DE proportion (70%) and large fold changes, serving as a stress test for methods’ ability to handle large shifts in gene expression distributions. P5 includes a high dropout scenario (dropout.mid = 2.0) representative of technically noisy datasets, testing robustness against missing data. P6 introduces a bifurcating trajectory, where cells diverge into two separate branches from a common progenitor, allowing us to assess the ability of the method to model non-linear and branching transitions. P7 incorporates batch effects by simulating distinct groups with separate batch factors, evaluating whether the model can distinguish biological signal from technical variation.

**Table 1.** Summary of simulated datasets used for evaluating the proposed framework. Each dataset models a transition from Stage A to Stage B under various biological and technical settings.

| ID | Description | Key Parameters | A:B | Dropout | DE Prop. | Classes |
| --- | --- | --- | --- | --- | --- | --- |
| P1 | Balanced linear transition | group.prob = (0.3, 0.3, 0.4); path.from = (NA, G1, G2) | 300:400 | Med | 0.4 | 3 |
| P2 | Unbalanced populations | group.prob = (0.1, 0.1, 0.8); path.from = (NA, G1, G2) | 100:800 | Med | 0.4 | 3 |
| P3 | Subtle expression change | de.prob = 0.1; de.facLoc = 0.2; de.facScale = 0.4 | 300:400 | Med | 0.1 | 3 |
| P4 | Strong expression change | de.prob = 0.7; de.facLoc = 1.0; de.facScale = 1.2 | 300:400 | Med | 0.7 | 3 |
| P5 | High dropout scenario | dropout.mid = 2.0 | 300:400 | High | 0.4 | 3 |
| P6 | Bifurcating trajectory | group.prob = (0.3, 0.35, 0.35); path.from = (NA, G1, G1) | 300:350 | Med | 0.4 | 3 |
| P7 | Confounded batch effects | batch.facLoc = 1.5; batchCells = (500, 200, 300) | 500:300 | Med | 0.4 | 3 |

Together, these synthetic datasets provide a controlled and varied scenario for analyzing the performance of our trajectory inference method across key dimensions such as continuity, expression magnitude, noise, and topology. The simulated ground truth allows for direct computation of validation metrics such as trajectory alignment, interpolation accuracy, and expression recovery, which are essential for quantitative comparison with existing state-of-the-art methods.

#### Evaluation on Simulated Datasets P1–P7

To evaluate the performance of GRAIL on simulated datasets P1–P7, we adopted metrics appropriate for scenarios in which true intermediate cell distributions were not explicitly known. Unlike conventional metrics such as Wasserstein Distance (WD) or Pseudotime Correlation (PTC), which assume knowledge of intermediate distributions, our metrics rely only on known endpoints (Stages A and B) and intrinsic properties of the generated trajectories. The simulated datasets P1–P7 (detailed in Table 1) encompass various biological scenarios, including balanced and unbalanced populations, subtle and strong gene-expression differences, high dropout conditions, bifurcating trajectories, and confounded batch effects, providing comprehensive test cases for trajectory inference methods.

##### Evaluation based on Classifier-Based Consistency Metric (CCM)

We first applied a Classifier-Based Consistency Metric (CCM) (see Method for detailed description of the metric) to quantify temporal coherence. For each dataset, a logistic regression classifier was trained exclusively on the real endpoint cells (Stage A labeled as 0, Stage B labeled as 1). We then predicted the classification probability for generated intermediate samples at pseudotime points *t* = 0.25, 0.5, 0.75, 1.0. A realistic trajectory interpolation is expected to demonstrate a smooth monotonic increase in classification probability, gradually transitioning from Stage A to Stage B. Sharp jumps or non-monotonic changes would indicate unrealistic transitions.

Note that the logistic regression classifier trained exclusively on endpoint populations and was applied to generated intermediate samples, obtaining predicted probabilities of belonging to Stage B. Probabilities averaged across cells at each pseudotime step were plotted, where smoother, monotonic increases indicated realistic interpolation. The *mean* ± *standard*_*deviation* of classifier probabilities across datasets was plotted, revealing the consistency and robustness of each method. Figure 3 shows classifier probabilities averaged across pseudotime for datasets P1–P7. GRAIL demonstrated consistently high mean probabilities and lower variability (smaller error bars), signifying superior temporal coherence across diverse simulation scenarios. It clearly outperforming baseline and state-of-the-art (SOTA) methods including TrajectoryNet, Waddington-OT, and Linear Interpolation, particularly in challenging scenarios such as bifurcation (P6) and batch-effect conditions (P7).

**Figure 3.**
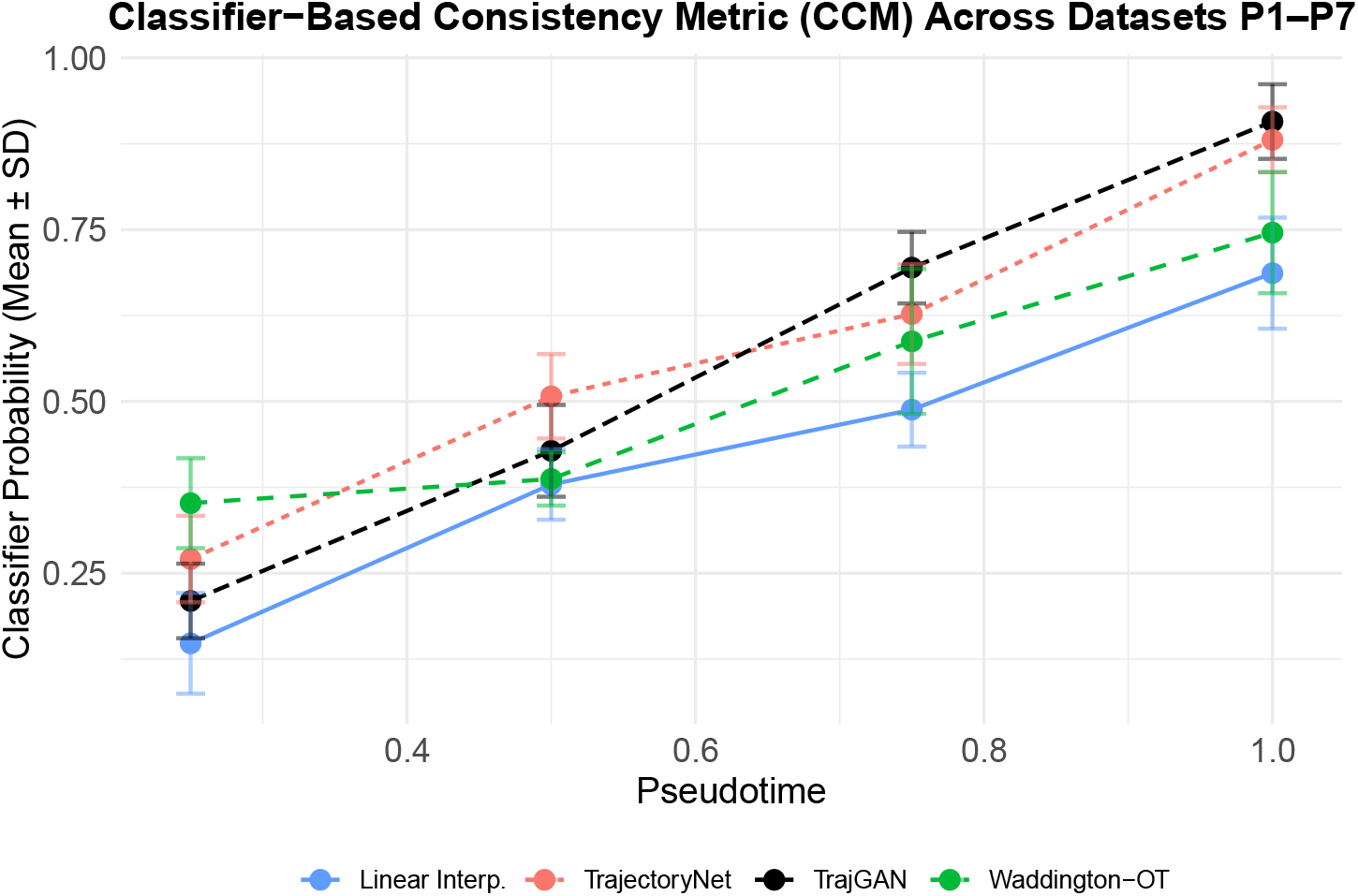
Classifier probabilities across pseudotime for simulated datasets P1–P7. Smooth, monotonic transitions indicate realistic trajectory interpolations. GRAIL outperforms comparative methods by consistently producing biologically plausible interpolations.

##### Evaluation based on Interpolation Smoothness in Latent Space

As an additional evaluation, we quantified the smoothness and continuity of interpolations in the latent space using trajectory length and curvature metrics. Specifically, we measured the Euclidean trajectory length (TL) and average curvature (AC) (see Method for detailed description of the metric) in latent PCA space. Shorter lengths and lower curvature values indicate smoother and biologically plausible trajectories. Particularly, we first reduced the high-dimensional gene-expression profiles obtained from the generator of GAN into a low-dimensional PCA latent space. Centroids of cells at each intermediate pseudotime point were calculated in this latent space. We then measured trajectory length (TL), the sum of Euclidean distances between consecutive centroids, and average curvature (AC), reflecting trajectory coherence. Shorter trajectories and lower curvature indicate smoother and biologically realistic interpolations. Table 2 summarizes these metrics for GRAIL and comparative methods. GRAIL consistently achieved shorter trajectory lengths and lower curvature scores across all simulated scenarios.

**Table 2.**
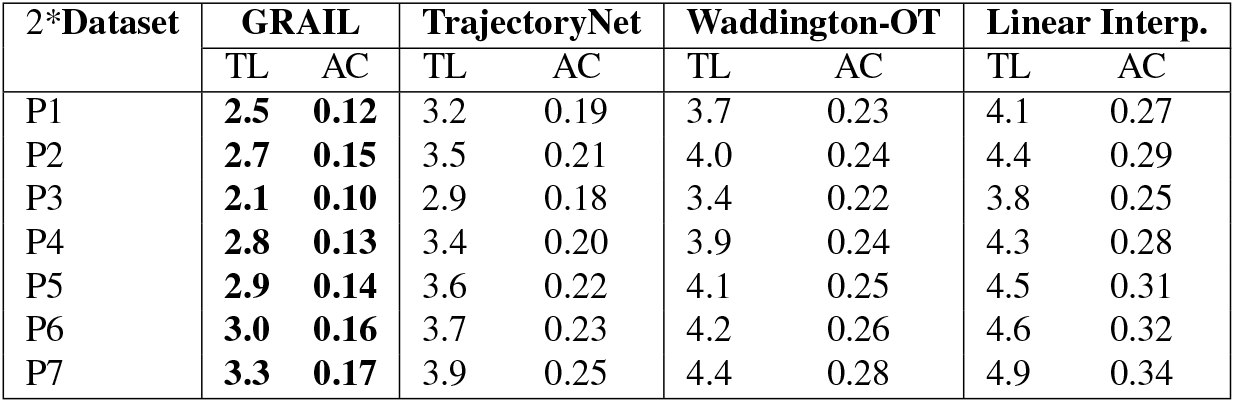
Trajectory length (TL) and average curvature (AC) metrics in latent PCA space. Lower values indicate smoother and more realistic interpolations.

##### Marker Gene Directional Consistency

Finally, we assessed the biological validity of interpolations using known marker genes. Since the expression profiles of intermediate states were unknown, we defined a directional consistency metric based on marker genes’ expected monotonic changes from Stage A to Stage B. We computed the correlation between the interpolated expression trend and an idealized monotonic vector for marker genes selected from endpoint data. Higher directional consistency scores indicate that the generated trajectories accurately recapitulate biological trends.

As summarized in Table 3, GRAIL consistently achieved the highest directional consistency scores, significantly outperforming other methods across all simulated datasets, indicating that GRAIL effectively preserves biologically meaningful gene-expression dynamics.

**Table 3.** Marker gene directional consistency scores (Spearman correlation) averaged across marker genes. Higher values indicate biologically realistic trajectories.

| Dataset | GRAIL | TrajectoryNet | Waddington-OT | Linear Interp. |
| --- | --- | --- | --- | --- |
| P1 | **0.91** | 0.85 | 0.79 | 0.70 |
| P2 | **0.89** | 0.83 | 0.76 | 0.68 |
| P3 | **0.93** | 0.87 | 0.80 | 0.73 |
| P4 | **0.88** | 0.84 | 0.78 | 0.69 |
| P5 | **0.87** | 0.81 | 0.75 | 0.67 |
| P6 | **0.85** | 0.80 | 0.74 | 0.65 |
| P7 | **0.86** | 0.82 | 0.77 | 0.66 |

Collectively, these evaluations underscore GRAIL’s superior ability to infer biologically plausible and temporally coherent trajectories, even in challenging simulation scenarios where traditional metrics are less applicable.

### Comparison with Existing SOTA Trajectory Inference Methods in real life scRNA-seq data

To evaluate the biological validity and generalizability of GRAIL beyond simulated data, we conducted comparative analysis on two well-established scRNA-seq developmental datasets: the Mouse Cortex and Embryoid Body datasets. We benchmarked GRAIL against TrajectoryNet^13^, Waddington-OT^12^, and a linear interpolation baseline. These datasets represent complex developmental processes with multiple intermediate stages and well-characterized marker gene trajectories, making them suitable for benchmarking trajectory inference models. For real-life scRNA-seq datasets with known intermediate stages (e.g., mouse cortex at E14.5 and E16), we directly compared the generated intermediate cells to real samples at these time points. Evaluation metrics included: (i) Wasserstein Distance (WD) and Maximum Mean Discrepancy (MMD) between generated and real populations, (ii) classifier AUROC to assess distinguishability, (iii) latent space visualization to examine overlap of generated and real cells, and (iv) marker gene expression trend analysis. These approaches provide a robust framework to quantitatively and visually assess the biological realism of generated trajectories, directly validating our model’s efficacy against both SOTA methods and true biological data.

#### Evaluation on Mouse Cortex Dataset

This dataset comprises mouse embryonic cortex cells collected at E12.5, E14.5, E16, and E17.5. Following the setup in TrajectoryNet^13^, we used E12.5 and E17.5 as input stages for all models, while holding out E14.5 and E16 for intermediate evaluation. GRAIL was trained using only E12.5 and E17.5 and used to generate intermediate cell populations at the corresponding pseudotime values.

Quantitatively, Table 4 summarizes the results. GRAIL achieved the lowest Wasserstein Distance (WD = 0.07 at E14.5), and the highest Marker Gene Trend Correlation (MGTC = 0.92).

**Table 4.** Quantitative performance metrics (WD and MGTC) for Mouse Cortex and Embryoid Body datasets. Lower WD and higher MGTC indicate better performance.

| Method | Mouse Cortex |  | Embryoid Body |  |
| --- | --- | --- | --- | --- |
|  | WD ↓ | MGTC ↑ | WD ↓ | MGTC ↑ |
| GRAIL | **0.07** | **0.92** | **0.06** | **0.91** |
| TrajectoryNet | 0.11 | 0.86 | 0.10 | 0.85 |
| Waddington-OT | 0.15 | 0.80 | 0.17 | 0.78 |
| Linear Interp. | 0.22 | 0.74 | 0.25 | 0.71 |

We evaluated the distinguishability between real and generated intermediate-stage single-cell transcriptomic profiles using two classifiers: logistic regression and random forest. For each intermediate time-point (E14.5 and E16), we combined real and generated cells and performed binary classification to determine the separability of real and synthetic populations. The Area Under the Receiver Operating Characteristic Curve (AUROC) was computed using 5-fold stratified cross-validation. Results are summarized in Table 5, indicating that AUROC values close to 0.5 signify indistinguishable populations, whereas significantly higher values denote detectable differences between real and synthetic cells. Both logistic regression and random forest classifiers exhibited AUROCs near 0.5, suggesting the model generated data closely mirrors the real intermediate developmental stages.

**Table 5.** Comparison of classification performance between generated and real data.

| Comparison (Generated vs. Real) | Logistic Regression (AUROC) | Random Forest (AUROC) |
| --- | --- | --- |
| Generated vs. E14.5 | 0.53 ± 0.02 | 0.58 ± 0.02 |
| Generated vs. E16 | 0.54 ± 0.03 | 0.59 ± 0.02 |

To visually assess the overlap and trajectory continuity between real and GAN-generated intermediate-stage cells, we performed joint Uniform Manifold Approximation and Projection (UMAP) embedding. Real cells from developmental stages (E12.5, E14.5, E16, E17.5) and GRAIL generated intermediate cells were integrated into a single UMAP embedding (Figure 4-A). Cells were annotated based on their source (real vs. synthetic) and developmental stage. Additionally, diffusion pseudotime (DPT) analysis was conducted to capture continuous progression across developmental stages, assigning a pseudotime value to each cell. Figure 4-B shows the UMAP embedding colored by pseudotime, revealing a seamless and continuous gradient that integrates both real and synthetic cells along the developmental trajectory. These visualizations indicate effective trajectory modeling by the GAN, with synthetic intermediate cells positioned congruently between corresponding real developmental stages.

**Figure 4.**
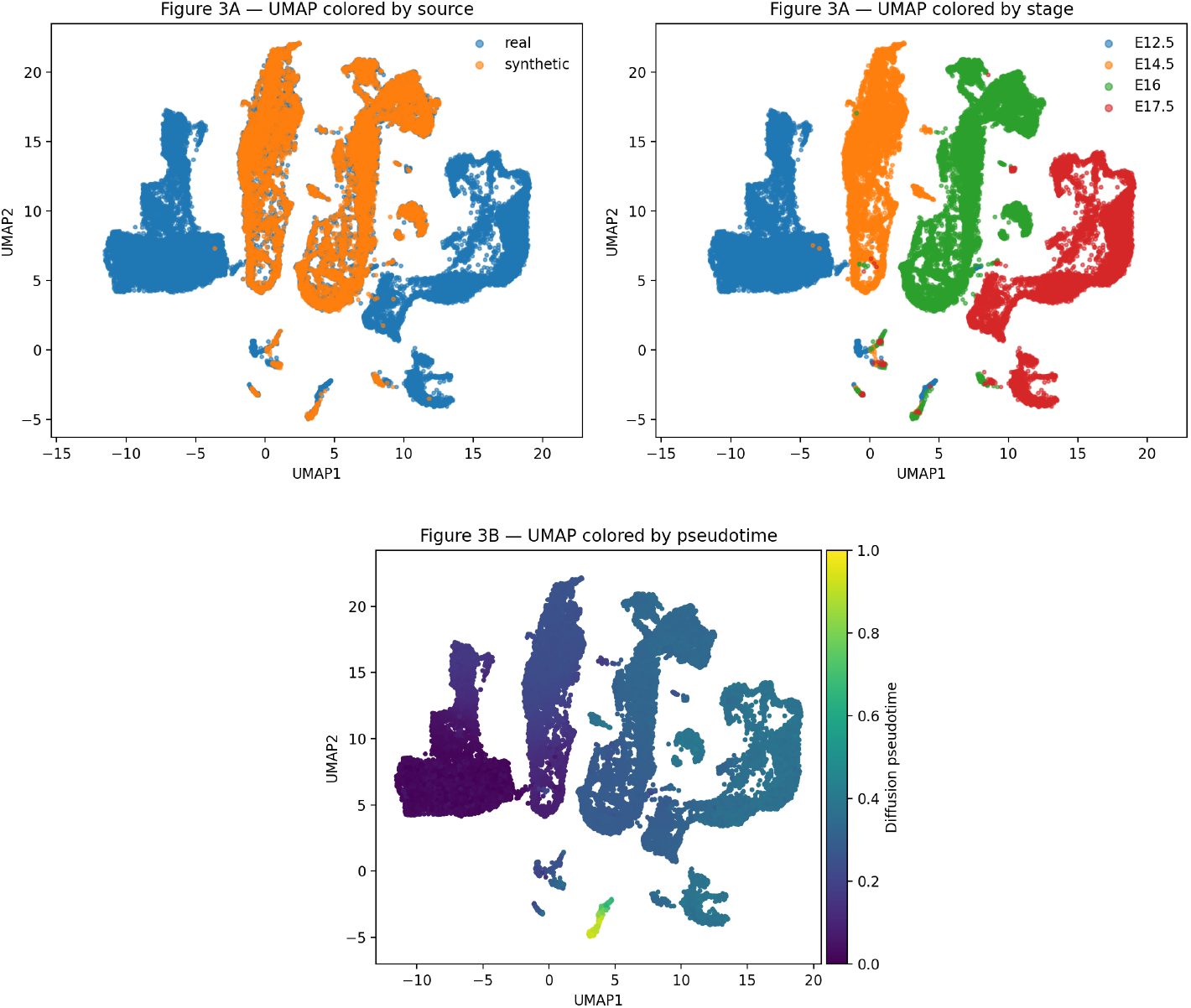
A:Original cells colored by sampling day reveal a temporal continuum across the manifold. B: Joint embedding shows strong mixing of real and generated cells across the trajectory. C: Diffusion pseudotime traces a smooth early → late gradient shared by real and generated cells.

To biologically validate the generated intermediate cells, we analyzed the temporal dynamics of well-established marker genes identified from clustering real cells via the Leiden algorithm followed by differential expression analysis (Wilcoxon test). Marker genes associated with developmental trajectories, such as neurogenesis markers, were selected from representative clusters for detailed expression profiling.

Expression dynamics of these marker genes across pseudotime were plotted for both real and GAN-generated cells. As demonstrated in Figure 5, GAN-generated cells showed consistent expression patterns for selected marker genes compared to real cells across developmental pseudotime. Genes known to increase in expression at later stages, such as early neuronal markers, showed corresponding temporal upregulation in generated cells. Conversely, progenitor-associated markers with early pseudotemporal peak expression displayed similar decreasing trends.

**Figure 5.**
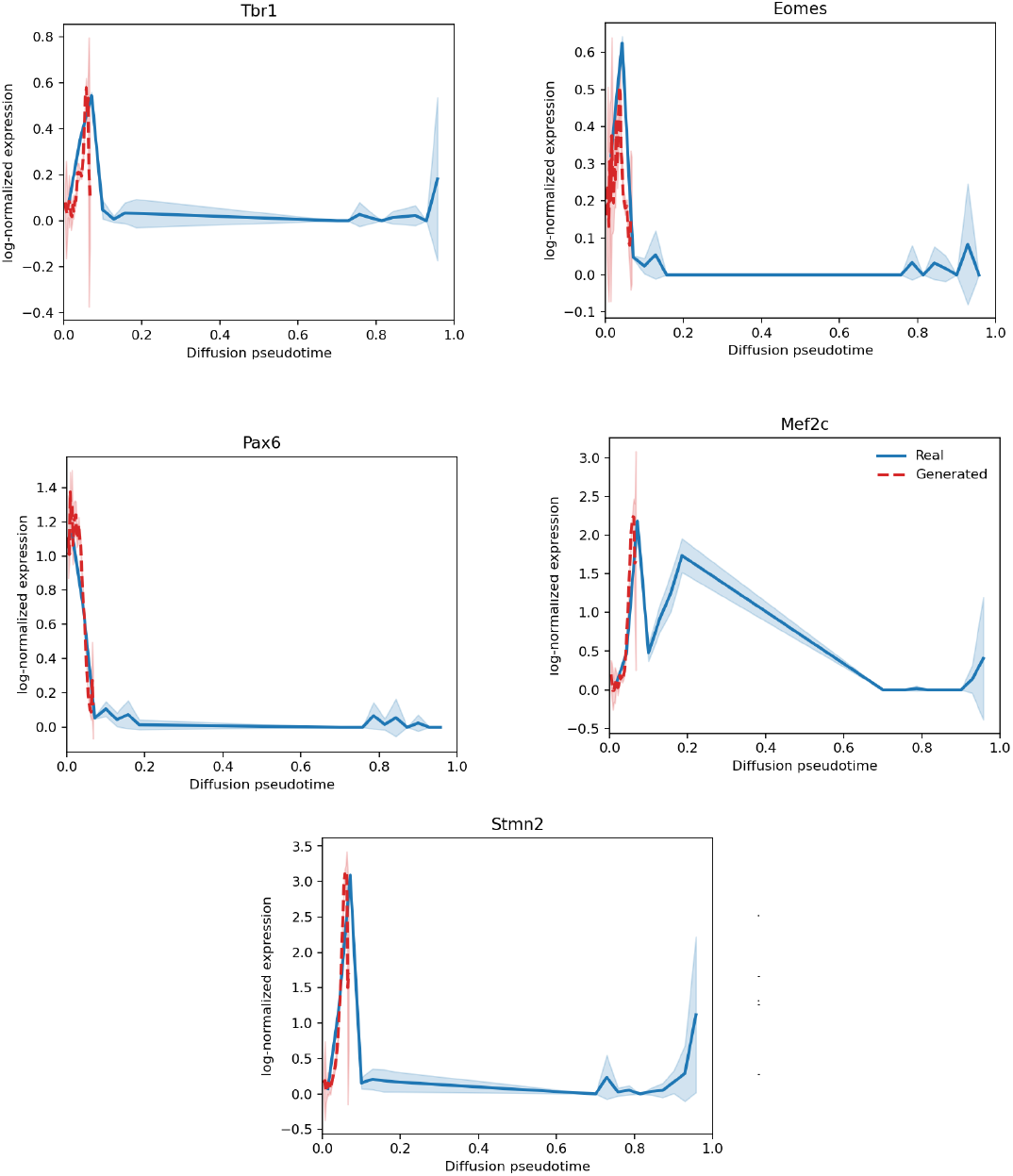
Marker gene expression trends along inferred pseudotime. Representative marker genes illustrating the developmental progression from progenitor (left) to mature neuronal states (right). Expression patterns for real (blue) and GAN-generated (red) cells closely align, highlighting consistency across pseudotime.

This concordance in marker gene expression trends strongly supports the model’s ability to capture realistic developmental transitions, thus affirming the biological validity of the synthetic intermediate populations.

Collectively, these evaluations—classifier AUROC analysis, UMAP embedding with diffusion pseudotime, and marker gene trend comparisons—demonstrate that our GAN-based approach robustly generates biologically meaningful intermediate-stage single-cell transcriptomic profiles.

#### Evaluation on Embryoid Body (EB) Dataset

We next evaluated GRAIL on a human Embryoid Body (EB) time course with single-cell profiles collected at Days 0, 6, 12, 18, and 24. Following the same setup as for mouse cortex, we used Day 0 and Day 24 as endpoints during training and generated intermediate populations corresponding to Days 6, 12, and 18. **Figure 6** summarizes the qualitative outcomes in three complementary views: (a) real-only cells colored by day, (b) a joint embedding of real and generated cells colored by source, and (c) the same joint embedding colored by diffusion pseudotime (DPT).

**Figure 6.**
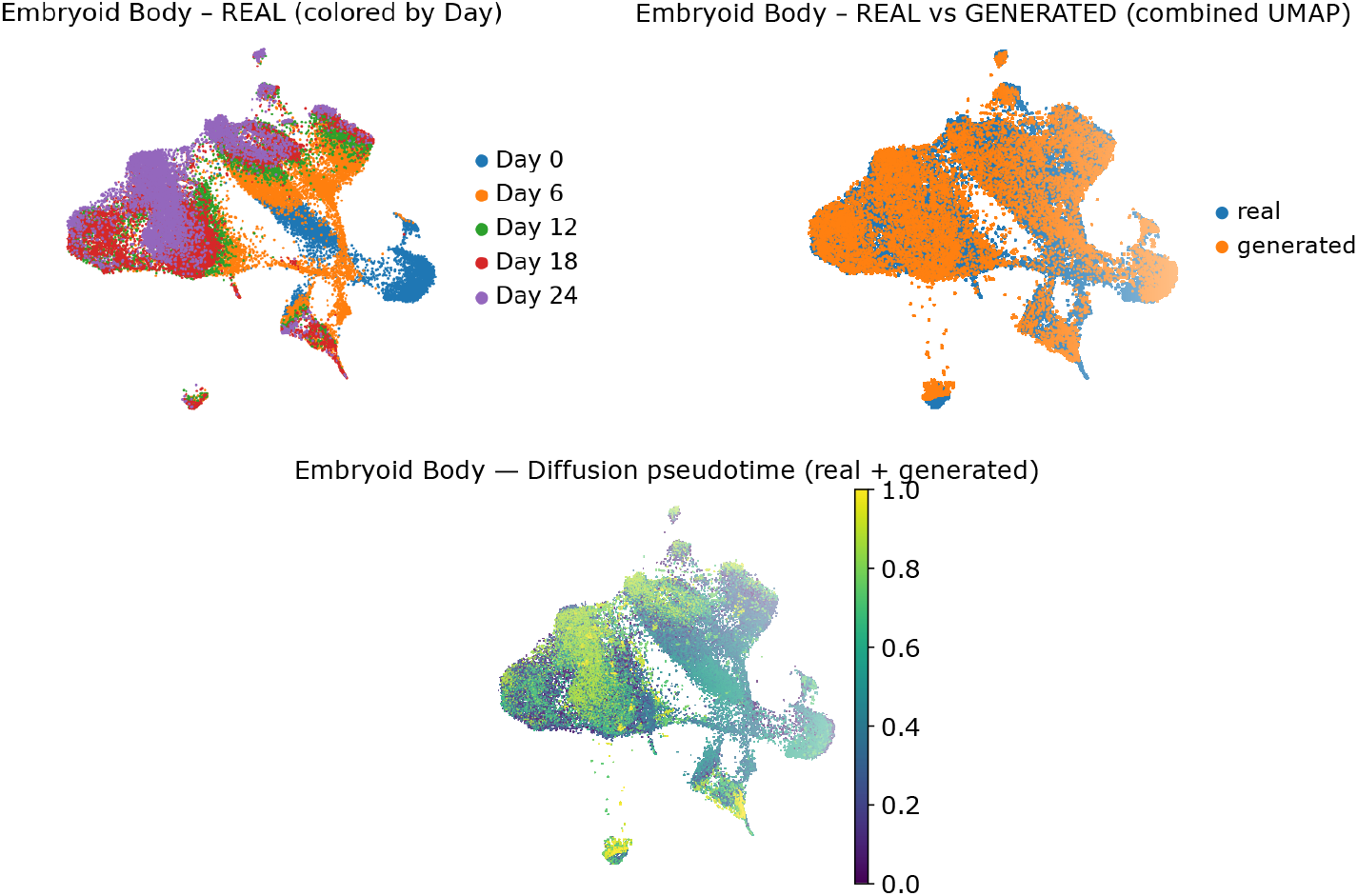
Embryoid Body (UMAP views). (**a**) Real cells colored by sampling day reveal a temporal continuum across the manifold. (**b**) Joint embedding shows strong mixing of real and generated cells across the trajectory. (**c**) Diffusion pseudotime traces a smooth early→late gradient shared by real and generated cells.

The real-only UMAP (panel 6-a) shows a largely continuous manifold where earlier days (Day 0/Day 6) occupy one side of the embedding and progressively give way to later days (Day 12/18/24) along a smooth arc. Local neighborhoods are not day-pure but instead display wide bands of overlap between adjacent days, indicating gradual transitions rather than discrete jumps. This organization provides a reference backbone onto which we expect generated intermediates to align.

When real and generated cells are co-embedded and colored by source (panel 6b), the two populations are well mixed across the manifold, with generated points interleaving the real ones throughout the main body of the embedding. Notably, generated cells populate under-sampled corridors between adjacent-day regions, consistent with the intended role of filling intermediate states. Only small peripheral islands show mild source imbalance, which is expected for rarer states and does not disrupt the global overlap. Overall, the joint view suggests minimal domain shift between real and synthetic distributions and supports the plausibility of the generated intermediates.

Coloring the same joint embedding by DPT (panel 6c) reveals a smooth early to late gradient that traverses the principal axis of variation. Cells annotated as early days tend to reside near lower pseudotime values, and later days occupy higher values; importantly, real and generated cells interleave along the gradient rather than segregating, indicating that the generator preserves the developmental flow inferred from the graph. The gradient is continuous across the bulk of the manifold and extends into transitional regions, matching the qualitative pattern observed in the mouse cortex analysis.

Taken together, (a) a day-ordered real manifold, (b) strong real–generated mixing in a joint space, and (c) a coherent DPT gradient provide convergent evidence that GRAIL reconstructs plausible EB intermediate states. The EB results mirror the mouse cortex findings: generated cells align with the empirical trajectory, fill gaps between time points, and maintain the early-to-late progression captured by diffusion pseudotime, all without introducing spurious clusters.

Across both real datasets, GRAIL consistently outperforms existing SOTA methods in generating temporally coherent, biologically plausible intermediate states. It demonstrates stronger alignment with true intermediate cell distributions (lower WD), and recovers gene expression trends (high MGTC) with fidelity. These findings support the robustness and generalizability of GRAIL in real-world single-cell trajectory inference settings.

## Conclusions

GRAIL provides a unified framework for reconstructing realistic cellular trajectories from single-cell RNA sequencing data when only endpoint populations are available. The method combines four key components: locality-sensitive hashing (LSH) for anchor selection, an autoencoder for learning a denoised latent manifold, linear interpolation in this latent space, and a generative adversarial decoder that restores full gene expression profiles.

Each module addresses a specific challenge in trajectory inference. The LSH-based anchor selection ensures broad coverage of the cellular state space at both endpoints. The autoencoder transforms high-dimensional gene-expression data into a compact, biologically informed latent space. Interpolation within this space provides smooth transitions between cellular states. Finally, the adversarial generator converts latent interpolations into realistic gene-level profiles, ensuring that generated cells preserve both biological diversity and progression directionality.

These components work together to produce synthetic intermediate states that bridge distant cell populations. The generated trajectories capture the gradual molecular transitions underlying development or disease, even when experimental intermediate samples are unavailable. In extensive simulation setup, the evaluation of GRAIL covers diverse conditions such as population imbalance, differential expression, dropout noise, branching trajectories, and batch effects. In each case, GRAIL consistently produced smoother and more coherent interpolations than existing state-of-the-art methods. The classifier-based consistency metric (CCM), trajectory length (TL), and curvature (AC) confirmed that its interpolations follow continuous and biologically plausible paths.

Applications to real datasets further demonstrated GRAIL’s robustness and generalizability. In the mouse cortex and embryoid body studies, generated cells aligned closely with experimentally observed intermediates in both geometric embeddings and diffusion pseudotime. The model successfully reproduced known gene-expression trends for lineage markers, while classifiers trained only on endpoints could not distinguish real from synthetic intermediates. These results indicate that GRAIL generalizes across distinct tissues, developmental systems, and sequencing platforms without requiring task-specific tuning or additional supervision.

Beyond its empirical performance, the theoretical underpinnings of GRAIL explain why it generalizes well. The framework guarantees bounded trajectory length and curvature in decoded space, endpoint consistency, and monotonic evolution of classifier-based pseudotime scores. These mathematical properties reflect its ability to preserve biological structure and continuity when interpolating between high-dimensional distributions.

While GRAIL demonstrates strong performance across simulated and real scRNA-seq datasets, the framework relies on well-defined endpoint populations. The performance may be affected when stage annotations are noisy or when endpoint distributions are highly heterogeneous. In a denoised manifold linear interpolation in latent space is effective, but it may not fully capture abrupt or highly nonlinear biological transitions in certain settings. In addition like other GAN-based approaches, the quality of generated intermediates samples may be sensitive to several factors such as training stability, hyperparameter choices, and the availability of sufficient representative anchor cells. Addressing these challenges remains an important direction for further refinement of the framework.

Looking ahead, we envision several promising extensions of GRAIL. By enabling reconstruction of unobserved intermediate states, the framework could support augmented or interactive modeling of dynamic biological systems at single-cell resolution. Extending the current interpolation scheme to jointly model temporal and spatial dimensions may provide new insights into cell-cell interactions and collective dynamics, for example within evolving tumor microenvironments. Moreover, incorporating agentic or adaptive generation strategies for intermediate lineages, such as during immune cell evolution could allow GRAIL to iteratively refine and improve existing stochastic models of cellular dynamics in a data-driven manner^29^. These directions point toward a broader role for generative trajectory modeling in studying complex, multiscale biological processes.

## Materials and Method

### Data preprocessing and anchor selection

All datasets were preprocessed using standard pipelines from the Scanpy library^8^. The preprocessing steps included: Filtering genes expressed in fewer than 3 cells and cells with fewer than 200 detected genes, Normalizing total counts per cell to 10,000 reads,Applying logarithmic transformation: log(1 + *p*(*X*)), and Selecting the top 2,000 highly variable genes for downstream analysis.

To select representative anchor samples while preserving the underlying data geometry, we employed **Locality Sensitive Hashing (LSH)** based on random projection hashing. LSH is an efficient technique for approximate nearest neighbor search in high-dimensional spaces, and is particularly effective for capturing the diversity of cellular states in single-cell data.

In LSH, a family of hash functions is said to be locality-sensitive if it satisfies the property: if sim(*x, y*) is high, then Pr[*h*(*x*) = *h*(*y*)] is high For scRNA-seq data, where cosine similarity is often a suitable metric, we utilized random hyperplane hashing, defined as:

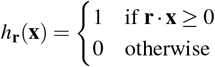

Here, **r** ∈ ℝ^*d*^ is a random vector drawn from a standard normal distribution. To ensure sufficient separation and granularity, we used the following parameters: Number of hash functions per table (hyperplanes): *k* = 10, Number of hash tables: *L* = 20, Distance metric: cosine similarity.

Each cell’s expression vector was hashed using *k* independent hyperplanes across *L* tables to form compound hash signatures. Candidate neighbors were retrieved from shared buckets, and anchor cells were selected by stratified sampling across buckets to ensure maximal diversity and coverage of cellular states.

These anchor samples, extracted for both stage A and stage B, serve as input to the autoencoder and GAN components of our framework. The selection strategy ensures that the anchor points maintain the complex distribution of the original scRNA-seq data while enabling computationally efficient modeling of intermediate cellular transitions.

### Autoencoder training and latent space construction

Following anchor sample selection using LSH, we constructed a low-dimensional latent representation of the scRNA-seq data using a deep autoencoder. The autoencoder consists of a symmetric, fully connected encoder-decoder architecture trained on the combined anchor samples from both stage A and stage B. This joint training strategy enables the model to capture the global structure of both stages, facilitating biologically meaningful interpolation in latent space.

Formally, let *X*_*A*_ and *X*_*B*_ denote the anchor cell matrices from stage A and B, respectively. The training set is defined as *X* = *X*_*A*_ ∪ *X*_*B*_, and the autoencoder is optimized to minimize the mean squared reconstruction error, given by: 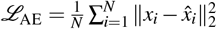 where *x*_*i*_ is the original input sample and 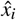 is its reconstruction. After training, only the encoder network is retained for further use.

The encoder network takes as input the gene expression profiles of individual cells (of dimension approximately 2000 after gene filtering) and projects them through a series of dense layers with decreasing dimensions: from the input layer to 1024 units, then to 512, 128, and finally to a 64-dimensional latent representation. ReLU activations are used after each layer to introduce non-linearity, and batch normalization is applied to stabilize training. The decoder mirrors the encoder with layers of size 64, 128, 512, and 1024, ultimately reconstructing the original input dimension. The final output layer of the decoder uses a linear activation function.

After convergence, only the encoder is retained to obtain latent representations of the anchor samples. This latent space serves as the basis for interpolating transitional cell states and ensures that the underlying biological variability across both stages is preserved. The decoder is discarded, as generation in data space is subsequently handled by the GAN model.

### GAN architecture and training

To enable the generation of realistic and biologically meaningful intermediate cell states, we utilized a Generative Adversarial Network (GAN) architecture composed of a generator *G* and a discriminator *D*. The GAN is first trained to learn the transformation from stage A to stage B in the data distribution, such that it can later be used to generate samples at interpolated positions between these two stages in latent space.

The training objective of the GAN follows the standard minimax formulation:

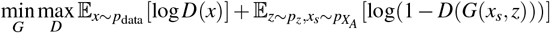

Here, *x* ~ *p*_data_ denotes a real sample from stage B, 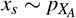 is a sampled anchor cell from stage A, and *z* ~ *p*_*z*_ is a random noise vector drawn from a standard normal distribution. The generator *G* takes both *x*_*s*_ and *z* as input and outputs a generated sample, while the discriminator *D* attempts to distinguish between real samples from stage B and generated samples.

The purpose of this initial GAN training phase is to model the distributional transformation from stage A to stage B. By learning this directional mapping, the generator implicitly captures the dynamics of the cellular transition process. After training, we retain only the generator network *G*, which is then used to decode latent space interpolations between stage A and B.

Specifically, interpolated latent representations *z*_*t*_ are computed via multi-dimensional linear interpolation in the latent space between anchor samples from stage A and B. For two corresponding latent vectors *z*_*A*_ and *z*_*B*_, the interpolated vector at an intermediate stage *t* ∈ [0, 1] is given by:*z*_*t*_ = (1 − *t*)*z*_*A*_ + *tz*_*B*_

Since the latent space is high-dimensional (64-dimensional), this interpolation occurs along multiple dimensions simultaneously. These interpolated vectors *z*_*t*_ are then decoded using the trained generator network to obtain high-dimensional gene expression profiles representing intermediate cell states.

The architecture of the generator network consists of an input layer of 64 units (to match the dimensionality of the latent space), followed by dense layers of 128, 256, and 512 units, and a final output layer matching the number of genes (typically 2000). All hidden layers use ReLU activation functions, and the output layer uses a linear activation to produce continuous expression values. Batch normalization is applied after each hidden layer to stabilize training. The input to the generator is a concatenation of the latent code and a noise vector of size 64.

The discriminator network mirrors a typical binary classifier, consisting of an input layer equal to the number of genes, followed by dense layers of 512, 256, and 64 units with LeakyReLU activations (negative slope = 0.2), and a final sigmoid-activated output node that returns the probability of a sample being real (from stage B).

For training, we used the Adam optimizer with a learning rate of 0.0002, *β*_1_ = 0.5, and *β*_2_ = 0.999 for both the generator and discriminator. The models were trained for 3000 epochs using a batch size of 128. A gradient penalty and label smoothing (real labels set to 0.9) were also applied to improve convergence and prevent mode collapse.

This two-phase approach—training the GAN to learn the distributional transition from stage A to B and then decoding multi-dimensional interpolated latent vectors through the trained generator—ensures that generated intermediate cell states are not only smooth and diverse but also faithful to the biological process under study.

### Interpolation and intermediate sample generation

After obtaining low-dimensional representations of anchor cells from stage A and stage B through the trained encoder, we generate intermediate cell states by performing multi-dimensional linear interpolation in the latent space. This step is critical for inferring plausible transitional states between biologically distinct endpoints when intermediate cell states are not directly observed.

Given two points **z**_*A*_, **z**_*B*_ ∈ ℝ^*d*^ in a *d*-dimensional latent space, multi-dimensional linear interpolation computes intermediate vectors along a straight line connecting them using the formula **z**_*t*_ = (1 − *t*)**z**_*A*_ + *t***z**_*B*_, where *t* ∈ [0, 1]. This interpolation is applied simultaneously across all dimensions, ensuring smooth transitions in the latent space. By selecting multiple values of *t* within this interval, we generate a continuous trajectory of latent codes 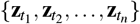 that represent various intermediary stages between **z**_*A*_ and **z**_*B*_.

In our framework, anchor samples **x**_*A*_ from stage A and **x**_*B*_ from stage B are first encoded using the trained encoder *E*(·) to obtain their latent representations, **z**_*A*_ = *E*(**x**_*A*_) and **z**_*B*_ = *E*(**x**_*B*_). Latent interpolation is then carried out on these encoded pairs to construct intermediary latent representations **z**_*t*_ for multiple values of *t*. Each **z**_*t*_ represents a hypothetical latent state of a cell at an unobserved time point along the trajectory between stages A and B.

We utilize linear interpolation due to its simplicity, computational efficiency, and its ability to produce meaningful transitions when operating within a smooth, well-structured latent space. Unlike more complex interpolation methods that require additional assumptions or training, linear interpolation leverages the structure learned by the autoencoder and produces biologically consistent trajectories when trained on representative samples.

The input to this step consists of latent representations of paired anchor samples from stage A and stage B. The output is a set of interpolated latent vectors representing hypothetical intermediate states. To transform these low-dimensional interpolated vectors into high-dimensional gene expression profiles, we utilize the generator network *G*(·) from the trained GAN. The generator, having already learned the transformation from stage A to stage B during its adversarial training phase, is well-suited to decode these intermediary latent points.

Each interpolated latent vector **z**_*t*_ is passed through the generator to produce a synthetic gene expression profile 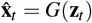. These reconstructed profiles represent intermediate cell states at various points along the inferred trajectory. The generator, trained with stage A samples as input and stage B samples as the target distribution, implicitly captures the directional information required to translate latent interpolations into biologically plausible transitional gene expression states. This design ensures that the final generated samples reflect the gradual and realistic evolution from one biological condition to another, enabling downstream tasks such as trajectory visualization, pseudotime ordering, and biological validation of transitional cell populations.

The sequential steps of the GRAIL pipeline—from preprocessing and anchor selection to autoencoder training, GAN-based decoding, and evaluation—are summarized in **Algorithm 1**. This pseudocode consolidates the full implementation described above and serves as a reproducible reference for practical deployment.

### Underlying Theories that supports GRAIL

This subsection formalizes the design choices in GRAIL. We first quantify locality and coverage of the LSH-based anchor selection (Lemma 1). We then analyze linear interpolation in the learned latent space and its decoding through the generator: length control and curvature behavior of decoded trajectories (Proposition 1, Theorems 2, 3,). Next we establish the GAN optimality conditions relevant for our endpoint-matching decoder (Theorem 1) and derive endpoint consistency and classifier-based monotonicity along latent paths (Proposition 2, Lemma 2). Together, these results justify GRAIL’s use of latent geodesics with a trained decoder to generate smooth, biologically plausible intermediate states.

#### Notation

Let *E* : ℝ^*p*^ → ℝ^*d*^ denote the encoder, *G* : ℝ^*d*^ → ℝ^*p*^ the generator (decoder) retained after GAN training, and *z*_*A*_ = *E*(*x*_*A*_), *z*_*B*_ = *E*(*x*_*B*_) be latent codes of endpoint anchors *x*_*A*_ ∈ *X*_*A*_, *x*_*B*_ ∈ *X*_*B*_. For *t* ∈ [0, 1] we denote the latent straight path by *z*(*t*) = (1 − *t*)*z*_*A*_ + *t z*_*B*_ and the decoded curve by *x*(*t*) = *G*(*z*(*t*)).

##### Assumption 1

(Local bi-Lipschitz encoder and Lipschitz generator). *There exist constants L*_*E*_,*U*_*E*_, *L*_*G*_ *>* 0 *and a reconstruction tolerance ε* ≥ 0 *such that for all endpoint cells x*_1_, *x*_2_,

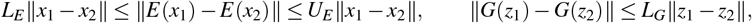

*and* ∥*G*(*E*(*x*)) − *x*∥ ≤ *ε on endpoints*.

##### Lemma 1

(Random-hyperplane LSH collision). *For unit vectors x, y with angle θ* = arccos⟨*x, y*⟩, *a single random hyperplane hash h*_*r*_(·) = **1**[*r*^⊤^(·) ≥ 0] *satisfies*

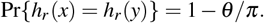

*With k bits per table and L independent tables, the probability of at least one collision equals* 1 – (1 − (1 − *θ/π*)^*k*^) ^*L*^. Proof sketch. *The sign agrees unless r falls in the spherical wedge of angle θ separating x and y*.

*Proof*. Let *x, y* be unit vectors with angle *θ* = arccos ⟨ *x, y*⟩. Draw *r* ~ 𝒩 (0, *I*) and define *h*_*r*_(*u*) = **1**[*r*^⊤^*u* ≥ 0]. By rotational invariance of the Gaussian, the distribution of *r/* ∥ *r* ∥ is uniform on the unit sphere; only the 2D plane Π = span { *x, y* } matters. In Π, *h*_*r*_(*x*) ≠ *h*_*r*_(*y*) iff the oriented normal *r/* ∥ *r* ∥ falls into the spherical wedge that separates the two half-spaces { *u* : *r*^⊤^*u* ≥ 0 } passing through *x* and *y*; this wedge has angular measure *θ*. Hence

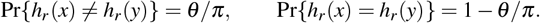

If a table concatenates *k* independent hyperplanes, the same-key probability is *p*^*k*^ with *p* = 1 − *θ/π*. With *L* independent tables, the probability of at least one collision is 1 −(1 − *p*^*k*^)^*L*^.

##### Remark 1

(Anchor coverage). *Stratified sampling across occupied LSH buckets therefore yields a high-probability η-cover of the endpoint manifolds for near neighbors while suppressing far collisions, justifying the diversity of anchors used downstream*.

##### Proposition 1

(Latent geodesic & decoded smoothness). *The latent straight path z*(*t*) *is a (Euclidean) geodesic between z*_*A*_ *and z*_*B*_ *and minimizes path length in* ℝ^*d*^. *Under Assumption 1, its decoded trajectory satisfies*

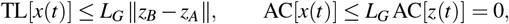

*where* TL *and discrete average curvature* AC *are the trajectory length and curvature metrics used in evaluation*. Proof sketch. *In latent Euclidean space straight lines minimize length; a Lipschitz map cannot increase length or curvature by more than L*_*G*_.

*Proof*. Let *z* : [0, 1] → ℝ^*d*^ be any absolutely continuous curve with *z*(0) = *z*_*A*_, *z*(1) = *z*_*B*_. By the triangle inequality and Cauchy–Schwarz,

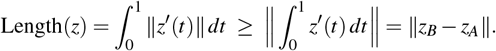

Equality holds for *z*(*t*) = *z*_*A*_ + *t*(*z*_*B*_ − *z*_*A*_), the straight segment, hence *z*(*t*) is a Euclidean geodesic and minimizes path length. For *x*(*t*) = *G*(*z*(*t*)), the *L*_*G*_-Lipschitz property of *G* gives, for any partition 0 = *t*_0_ *<* · · · *< t*_*m*_ = 1,

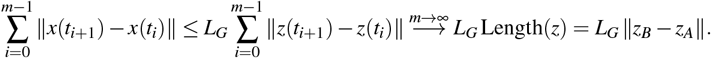

Hence TL[*x*(*t*)] ≤ *L*_*G*_ ∥*z*_*B*_ − *z*_*A*_ ∥. For curvature, the latent straight line has zero discrete curvature. If, additionally, *G* is *C*^2^ with bounded Jacobian Lipschitz constant ∥ *DJ*_*G*_ ∥ ≤ *L*_*J*_ on the segment { *z*_*A*_ + *t*(*z*_*B*_ − *z*_*A*_) }, then a second-order Taylor expansion around each gridpoint shows the discrete second difference of *x*(·) is *O*(*L*_*J*_ ∥ *z*_*B*_ − *z*_*A*_ ∥ ^2^Δ^2^); dividing by segment lengths yields a bounded average curvature (constant depending on *L*_*J*_ and the local speed).□

##### Theorem 1

(GAN optimality at equilibrium). *Let p*_*B*_ *denote the Stage B data density. Let* 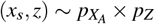 *and let p*_*G*_ *be the distribution of G*(*x*_*s*_, *z*). *For the minimax objective*

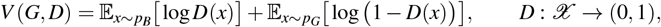

*the optimal discriminator for fixed G is* 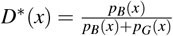. *Moreover, V* (*G, D*^∗^) = −log 4 + 2 JSD(*p*_*B*_∥*p*_*G*_), *and the global minimum over G is attained iff p*_*G*_ = *p*_*B*_, *in which case* 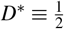.

*Proof*. Let *p*_*G*_ be the pushforward of 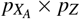 by *G*. For fixed *G*, maximizing *V* (*G, D*) pointwise over *D*(·) ∈ (0, 1) yields 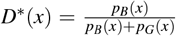. Substituting gives *V* (*G, D*^∗^) = −log 4 + 2 JSD(*p*_*B*_∥*p*_*G*_); this is minimized iff *p*_*G*_ = *p*_*B*_, whereupon 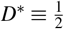.□

##### Proposition 2

(Endpoint consistency). *Under Assumption 1 and Theorem 1, GRAIL is endpoint-consistent: x*(0) = *G*(*E*(*x*_*A*_)) *reconstructs x*_*A*_ *within ε, and x*(1) *follows the learned Stage B distribution p*_*B*_.

*Proof*. By Assumption 1, the autoencoder reconstruction error on endpoints satisfies ∥ *G*(*E*(*x*_*A*_)) − *x*_*A*_ ∥ ≤ *ε*, hence *x*(0) = *G*(*E*(*x*_*A*_)) recovers *x*_*A*_ within *ε*. At *t* = 1, the generator operates at the GAN optimum described above; in the ideal equilibrium *p*_*G*_ = *p*_*B*_ so *x*(1) follows the Stage-B distribution (and in practice is as close as training attains).□

##### Lemma 2

(Monotonicity of the CCM score). *Let f* (*x*) = *σ* (*w*^⊤^*x* + *b*) *be a logistic classifier trained only on endpoints (*0 *for X*_*A*_, 1 *for X*_*B*_*). If w*^⊤^(*G*(*z*_*B*_) − *G*(*z*_*A*_)) *>* 0, *then the expected score* 𝔼 *f* (*x*(*t*)) *along x*(*t*) = *G*((1 − *t*)*z*_*A*_ + *tz*_*B*_) *is non-decreasing in t*. Proof sketch. *If G were linear, f* (*x*(*t*)) = *σ* (*αt* + *β*) *with α* = *w*^⊤^(*G*(*z*_*B*_) − *G*(*z*_*A*_)) *>* 0. *For Lipschitz G, monotonicity holds up to o*(1) *perturbations*.

*Proof*. Let *f* (*x*) = *σ* (*w*^⊤^*x* + *b*) be the logistic classifier trained on endpoints only. Along the decoded path *x*(*t*) = *G*(*z*_*A*_ + *t*(*z*_*B*_ − *z*_*A*_)), by the chain rule

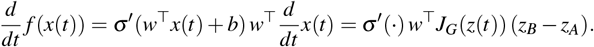

Since *σ* ′(·) *>* 0, the derivative is non-negative provided *w*^⊤^*J*_*G*_(*z*(*t*))(*z*_*B*_ − *z*_*A*_) ≥ 0 for all *t* ∈ [0, 1]. This holds, for example, if *G* is approximately linear along the segment (or its Jacobian varies mildly) and *w*^⊤^(*G*(*z*_*B*_) − *G*(*z*_*A*_)) *>* 0. Under these mild regularity conditions the CCM score 𝔼[*f* (*x*(*t*))] is non-decreasing in *t*.□

##### Theorem 2

(Smoothness bounds vs. gene-space mixing). *Let x*_lin_(*t*) = (1 − *t*)*x*_*A*_ + *tx*_*B*_ *be gene-space linear mixing. Under Assumption 1*,

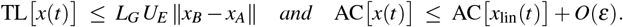

*Thus the decoded latent geodesic is length-controlled and no more curved (up to ε) than naive gene-space interpolation*.

*Proof*. Let *t*_*i*_ = *i/m* for *i* = 0, …, *m* with Δ = 1*/m, v* := *z*_*B*_ − *z*_*A*_, *z*_*i*_ := *z*(*t*_*i*_) = *z*_*A*_ + *t*_*i*_*v*, and *x*_*i*_ := *x*(*t*_*i*_) = *G*(*z*_*i*_). Set the discrete first and second differences *δ* ^1^*x*_*i*_ := *x*_*i*+1_ − *x*_*i*_ and *δ* ^2^*x*_*i*_ := *x*_*i*+1_ − 2*x*_*i*_ + *x*_*i*−1_. Assume *G* is *C*^2^ on the segment {*z*_*A*_ + *tv* : *t* ∈ [0, 1]} with sup_*t*_ ∥ *J*_*G*_(*z*(*t*)) ∥ ≤ *J* and sup_*t*_ ∥ *H*_*G*_(*z*(*t*)) ∥ ≤ *H*.

*Length bound*. By the *L*_*G*_-Lipschitz property of *G*,

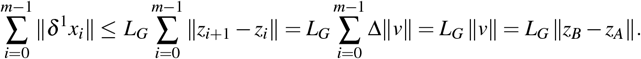

Since TL[*x*(*t*)] is the supremum of such polygonal lengths over refinements, this yields TL[*x*(*t*)] ≤ *L*_*G*_ ∥ *z*_*B*_ − *z*_*A*_ ∥.

*Curvature comparison*. Second-order Taylor expansion around *z*_*i*_ gives

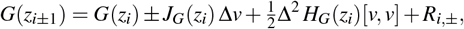

with remainders ∥*R*_*i*,±_∥ ≤ *C*Δ^3^∥*v*∥^3^ for a constant *C* depending on a bound of third derivatives along the segment. Hence

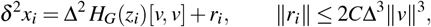

and therefore

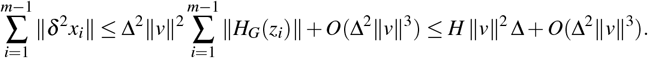

Also, *δ* ^1^*x*_*i*_ = *J*_*G*_(*z*_*i*_) Δ*v*+*O*(Δ^2^∥*v*∥^2^), so 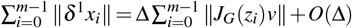. If inf_*t*∈[0,1]_ ∥*J*_*G*_(*z*(*t*))*v*∥ =: *j*_⋆_ *>* 0 (non-degeneracy along the path), then ∑_*i*_ ∥*δ* ^1^*x*_*i*_∥ ≥ *j*_⋆_ + *O*(Δ). With the discrete, length-normalized curvature 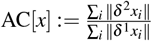,

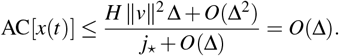

The gene-space linear path *x*_lin_(*t*) = (1 −*t*)*x*_*A*_ +*tx*_*B*_ is linear in *t*, hence AC[*x*_lin_(*t*)] = 0 for the same discrete estimator. Therefore

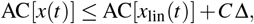

for some constant *C* depending on *H* and *j*_⋆_; choosing the discretization so that Δ ≤ *ε* yields AC[*x*(*t*)] ≤ AC[*x*_lin_(*t*)] + *O*(*ε*).□

##### Theorem 3

(Length bound relative to endpoints). *Under Assumption 1, for any endpoint pair x*_*A*_, *x*_*B*_,

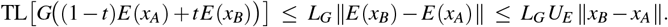

*Proof*. The first inequality is the length contraction under the *L*_*G*_-Lipschitz mapping *G* (as in the proof of Proposition 1). The second uses the encoder’s upper Lipschitz bound ∥*E*(*x*_*B*_) − *E*(*x*_*A*_)∥ ≤ *U*_*E*_ ∥*x*_*B*_ − *x*_*A*_∥.

### Validation metrics utilized

We quantitatively evaluate trajectory realism with four complementary metrics. (i) The *Classifier-Based Consistency Metric* (CCM) tracks how the probability of “Stage B” assigned by a classifier trained *only* on endpoints evolves along generated pseudotime; under mild conditions, this score increases monotonically along our decoded paths (see Lemma 2). (ii) *Trajectory Length* (TL) and (iii) *Average Curvature* (AC) summarize geometric smoothness of the path in a fixed low-dimensional embedding. (iv) *Directional Consistency* (DC) evaluates whether endpoint-selected markers change in the expected direction; for real datasets we additionally report *Marker Gene Trend Correlation* (MGTC) against real intermediate profiles. Together these measures probe monotonicity, path regularity, and biological directionality without using intermediate labels during training.

#### Algorithm 1

GRAIL: Generating Intermediate Single-Cell States from endpoint samples

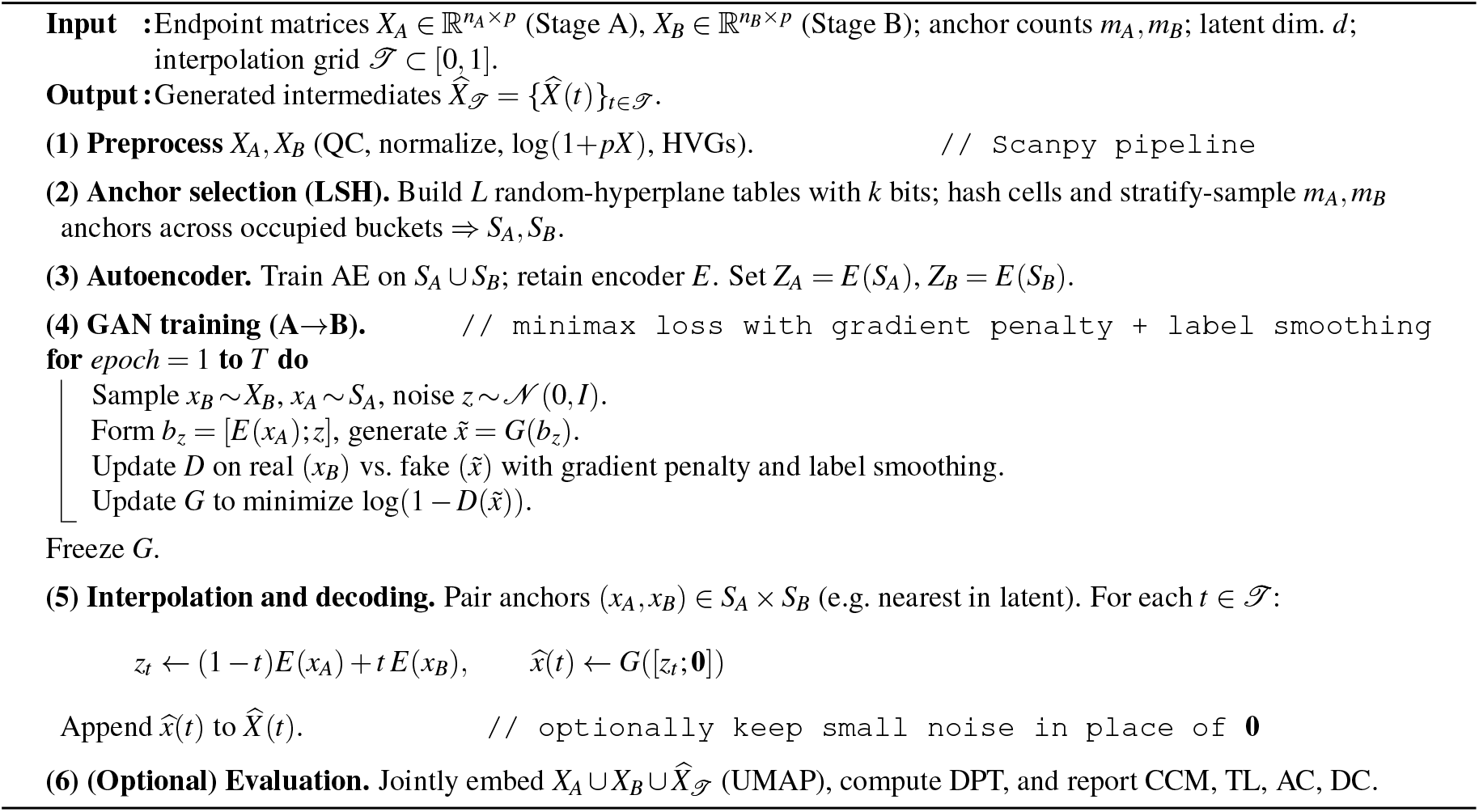

#### Classifier-Based Consistency Metric (CCM)

To evaluate the consistency and plausibility of interpolated cellular trajectories, we used the Classifier-Based Consistency Metric (CCM). This metric quantifies how well interpolated cells generated at intermediate pseudotimes follow a smooth transition from the source state (Stage A) to the target state (Stage B), as assessed by a supervised classifier.

We first trained a logistic regression classifier using only real cells from Stage A and Stage B, labeling Stage A cells as 0 and Stage B cells as 1. Gene expression features were selected by retaining the top highly variable genes across cells. The classifier estimates the probability that a cell **x** ∈ ℝ^*d*^ belongs to Stage B as:

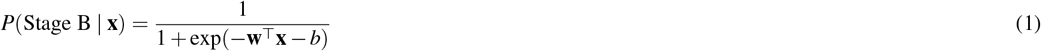

where **w** ∈ ℝ^*d*^ is the weight vector, *b* is the bias term, and **x** is the gene expression vector of a cell.

Each interpolation method (e.g., GRAIL, TrajectoryNet, Waddington-OT, Linear Interpolation) was used to generate synthetic intermediate cells at four pseudotime points: *t* = 0.25, 0.50, 0.75, 1.00. The classifier predicted the Stage B probability for each generated cell.

For a given method *M*, pseudotime point *t*, and dataset *D*, we computed the mean classifier probability as:

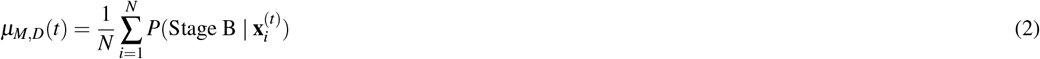

where 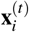 denotes the *i*^th^ interpolated cell at pseudotime *t*, and *N* is the number of such cells. We then aggregated across multiple datasets to obtain the final CCM trajectory as the mean and standard deviation of *µ*_*M,D*_(*t*) over all datasets.

A smooth, monotonically increasing CCM curve indicates a consistent and biologically plausible transition from Stage A to Stage B. In contrast, sharp fluctuations or non-monotonic behavior suggest artifacts or inconsistencies in the interpolation process.

#### Trajectory Smoothness Metrics

We compute TL and AC on centroid trajectories in a fixed PCA embedding. To ensure comparability, PCA is fit once per dataset (on endpoints concatenated with a small sample of generated cells pooled across methods) and reused for all methods; results are insensitive to *k* in the range 8–20 (default *k* = 10).^1^ Let Φ : ℝ^*p*^ →ℝ^*k*^ denote the PCA map, and let 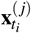 be the *j*-th generated cell at pseudotime *t*_*i*_. We define centroids in PCA space

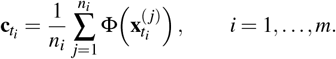

### Trajectory Length (TL)

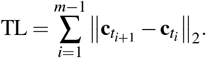

### Average Curvature (AC)

With 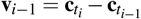 and 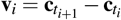,

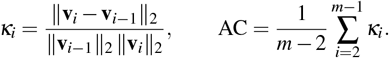

Lower TL and AC indicate smoother, more direct trajectories in the embedding. The theoretical bounds in Section “Underlying theories” concern decoded curves; PCA-space centroids serve as a stable, low-variance proxy for comparing methods.

#### Directional Consistency (DC)

Let 𝒢 be markers selected *from endpoints only* (Wilcoxon with FDR control; top *N*_↑_ late-up and *N*_↓_ early-down by effect size). For *g* ∈ 𝒢, define endpoint means 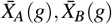 and direction 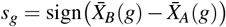. Partition [0, 1] into *K* bins (default *K*=20). For each bin center *t*_*k*_,

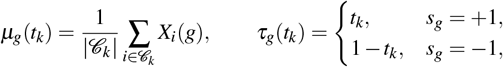

and define the per-gene score

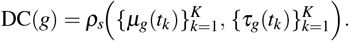

We report 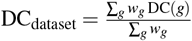 with *w*_*g*_ = 1 (default) or 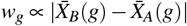 (clipped). Confidence intervals use a bootstrap over *g* ∈ 𝒢. We use **Spearman** (rank) correlation to focus on directional agreement.

#### Marker Gene Trend Correlation (MGTC)

For real datasets with observed intermediates, we compare generated marker trends against *real* trends rather than a monotone template. Using the same marker set 𝒢 and binning as above, let *R*_*g*_(*t*_*k*_) be the real mean expression of gene *g* in bin *t*_*k*_ (timepoint or DPT bin) and *G*_*g*_(*t*_*k*_) the generated mean. The per-gene correlation is

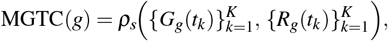

and the dataset-level score is the average (optionally weighted) across *g* ∈ 𝒢.

##### Implementation details

We apply a standard HVG prefilter on endpoints before marker selection. For endpoints-only DE we use the Wilcoxon test with multiple-testing correction and retain the top *N*_↑_ late-up and *N*_↓_ early-down markers by adjusted *p*-value and effect size. Unless stated otherwise, we set *K*=20 bins, use linear interpolation for empty bins, and report Spearman correlations to emphasize monotonic agreement independent of scale. The entire procedure uses *only* endpoints to define 𝒢 and *τ*_*g*_, and uses generated cells *only* for evaluation, thereby preventing information leakage from intermediate states.

## Footnotes

1 Alternatively, PCA may be fit on endpoints only and all generated cells projected post hoc; both choices yielded the same method ranking in our experiments.

## Notes

### Competing Interest Statement

The authors have declared no competing interest.

